# Neuronal subtype-specific metabolic changes in neurodegenerative and neuropsychiatric diseases predicted via a systems biology-based approach

**DOI:** 10.1101/2025.11.03.686281

**Authors:** Boyu Jiang, Shuang Wang, Junkai Xie, Hyunjin Kim, Anke M. Tukker, Juexin Wang, Aaron B. Bowman, Chongli Yuan, Priyanka Baloni

## Abstract

Understanding how distinct neuronal subtypes contribute to Alzheimer’s disease (AD) pathology remains a major challenge. Patient-derived induced pluripotent stem cell (iPSC) studies have shown neuronal subtype-specific molecular and pathological signatures, yet the underlying metabolic shifts driving this selective vulnerability are not completely understood. Here we present iNeuron-GEM, the first manually curated, genome-scale metabolic network of human neurons that integrates transcriptomic and metabolic knowledge to resolve subtype-specific metabolic states. By coupling iNeuron-GEM with single nucleus RNA sequencing data from post-mortem human cohort studies, ROSMAP and SEA-AD, we capture neuronal subtype-specific metabolic features and fluxes and identify perturbations in lipid and energy metabolism across excitatory and inhibitory neurons. Integrative analysis with NPS-AD data shows overlapping metabolic disruptions in AD and schizophrenia (SCZ), suggesting shared molecular vulnerabilities between neurodegenerative and neuropsychiatric disorders. We also developed a computational pipeline to infer transcriptional regulation of metabolic pathways and identify *NR6A1* and *NR3C1* as important regulators of lipid dysregulation in AD neurons. Our study establishes iNeuron-GEM as a framework to identify neuronal subtype-specific metabolic vulnerabilities in complex brain disorders.

## Introduction

The human brain has unique metabolic characteristics, including high energy demand and neurotransmitter cycling, both essential for maintaining normal neuronal function. During the progression of neurodegenerative diseases such as Alzheimer’s disease (AD), metabolic alterations in the human brain have been increasingly highlighted^1^. Positron emission tomography (PET) studies have found reduced glycolytic activity, an important process for ATP generation from glucose, in the brain of AD patients compared to healthy individuals^2^. Glycolytic dysfunction correlated with hallmark AD pathologies including amyloid-β (Aβ) deposition and tau hyperphosphorylation^3^. In addition to glucose metabolism, clinical evidence also demonstrated altered brain lipid metabolism in AD pathogenesis. Lipids, including fatty acids, cholesterol, glycerophospholipids, and sphingolipids, serve as both energy substrates and structural components. Abnormal lipid synthesis, degradation, and transport can promote Aβ aggregation, tau phosphorylation and synaptic functions, potentially contributing to AD pathogenesis^4,5^. Astrocyte-derived cholesterol has been shown to increase neuronal membrane cholesterol via apolipoprotein E (ApoE), a well-known genetic risk factor for AD^5,6^, thereby facilitating interactions between amyloid precursor protein (APP) and β- and γ-secretases, promoting Aβ formation^7^.

To understand mechanisms underlying such metabolic alterations, it is important to characterize metabolic activities of specific brain cell types. The brain is primarily composed of neurons, including excitatory and inhibitory neurons, and glial cells, including astrocytes, microglia, and oligodendrocytes. Post-mortem human brain single nucleus RNA-sequencing (snRNA-seq) studies have highlighted brain cell type-specific gene expression changes and metabolic shifts in AD^8,9^. For example, induced neurons derived from AD patient fibroblasts were reported to have a Warburg-like metabolic shift, exhibited by increase in aerobic glycolysis^10^, whereas ApoE4-mediated lipid dysregulation in microglia decreased Aβ uptake in the human induced pluripotent stem cell (iPSC)-derived models^11^.

Despite the abundance of postmortem multi-omics data from AD and healthy brains^8,12^, the lack of metabolomic resolution at the cell type level limits our understanding of neuron-specific metabolic reprograming in brain disorders. Computational modeling offers a powerful strategy to address this gap. Genome-scale metabolic models (GEMs) have the capability to integrate omics data with biochemical network knowledge to simulate cell type-specific metabolic states. GEMs are mathematical representations of biochemical networks which contain information of metabolic genes, enzymes/transporters, chemical reactions and metabolites^13^. By employing different constraint-based analyses, these models can simulate *in silico* metabolic flux distributions at a steady state^14^. Using systems biology approaches and integrative genome-scale metabolic models (GEM), we can identify metabolic shifts in different conditions. GEMs have already been widely applied to optimize metabolic engineering of bacteria and characterize metabolic alterations in human diseases, demonstrating advantages in exploring the relationship between genotypes and phenotypes^15,16^.

In this study, we report the first curated human neuronal subtype-specific GEM, namely *iNeuron-GEM*, constructed using snRNA-seq data from well-known AD cohort studies, namely the Religious Orders Study and Memory and Aging Project (ROSMAP) Study and the Seattle Alzheimer’s Disease Brain Atlas (SEA-AD) Study^8,12^. iNeuron-GEM simulates metabolic fluxes in excitatory and inhibitory neurons, enabling systematic identification of subtype-specific metabolic alterations in AD. To extend our findings, we integrated snRNA-seq data from the Mount Sinai Neuropsychiatric Symptoms in AD (NPS-AD) Study^17^, which includes individuals with AD, schizophrenia (SCZ), and mood disorders. Samples in NPS-AD cohort had information of the Braak stages (a widely applied measure of severity of neurofibrillary tangle (NFT) pathology^18^), that allowed us to investigate shared and distinct metabolic signatures across neurodegenerative and neuropsychiatric conditions. Our framework provides a systems-level platform for dissecting neuronal subtype-specific metabolic vulnerabilities and identifying potential regulatory mechanisms underlying AD progression.

## Results

### Construction of manually curated human neuron metabolic network – *iNeuron-GEM*

Using transcriptomics and metabolomics data, we generated the first curated human neuron-specific metabolic network, using Recon3D^19^, a generic human whole body metabolic network as the template (Figure 1A I). The *iNeuron-GEM* contained 1341 metabolite species, 3900 metabolic reactions and 1703 genes (Supplementary File 1 Table S1a, S1b, S1c). We used an iterative approach for generating the curated *iNeuron-GEM* metabolic network. We used the snRNA-seq data of postmortem prefrontal cortex samples from two AD cohorts, ROSMAP and SEA-AD studies, which contain information of 1032 individuals with an AD diagnosis (see Methods, Supplementary File 1 Table S2). To ensure specificity of our metabolic network, the draft reconstruction was further refined with untargeted metabolomics of human induced pluripotent stem cell (iPSC)-derived neurons and cerebrospinal fluid (CSF) (Table 1). We observed high overlap between experimental data and the metabolic genes and metabolites in the *iNeuron-GEM*, indicating cell type specificity. Our metabolic network was flux consistent (based on Memote analysis). 92% of metabolic genes in the metabolic network mapped with neuron-specific gene expression reported in the Human Protein Atlas (HPA) database ^20^. For intracellular metabolites in the metabolic network, about 55% of them had defined Human Metabolome Database (HMDB) ^21^ IDs and matched the metabolomics data from iPSC-derived neurons. 51% of the extracellular metabolites in the metabolic network were present in metabolomics data of CSF. As the synaptic region connects the neurons, we included a new compartment named “syna” to our metabolic network. This ‘synapse’ compartment included enzymes and transporter genes reported to be active in the synaptic region ^22^. It consisted of 36 reactions, most of which are involved in neurotransmitters transport (Supplementary File 1 Table S1a).

**Table 1.**
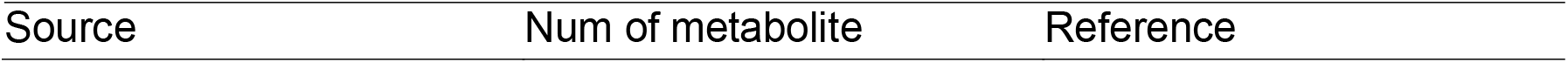
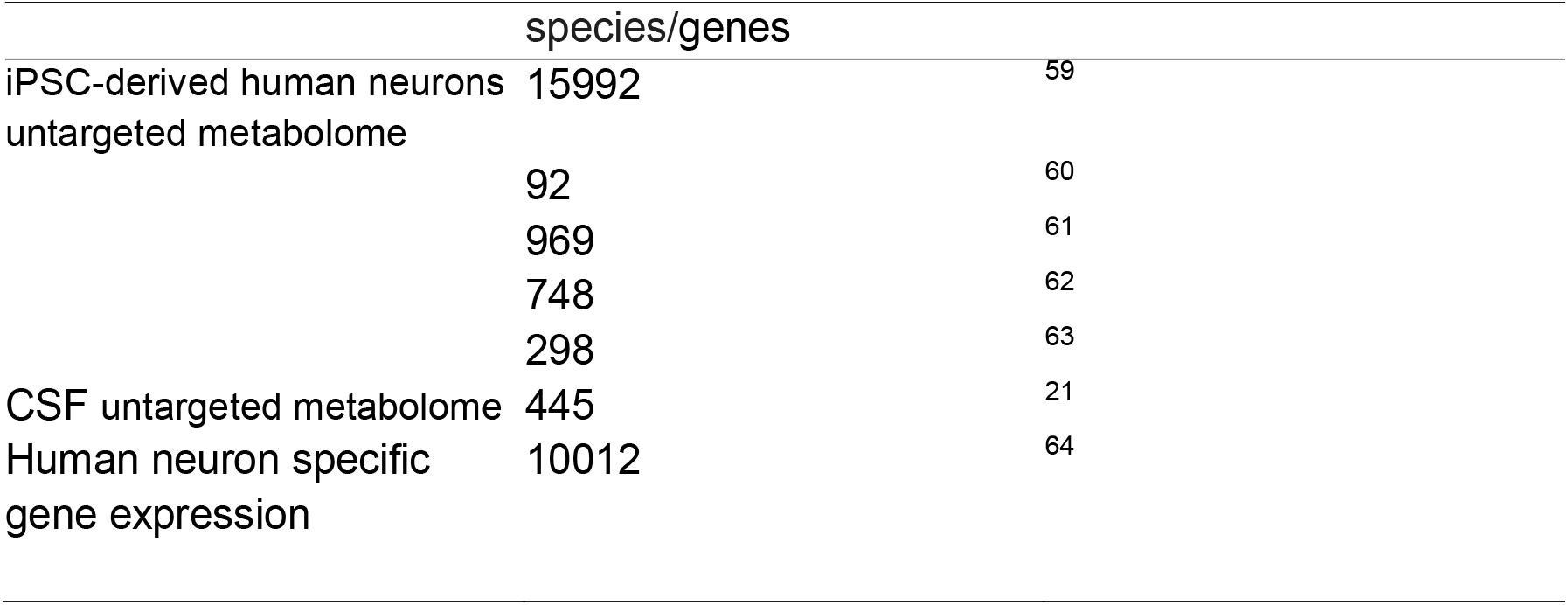
metabolomics and transcriptomics data used to curate human neuron reconstruction.

**Figure 1.**
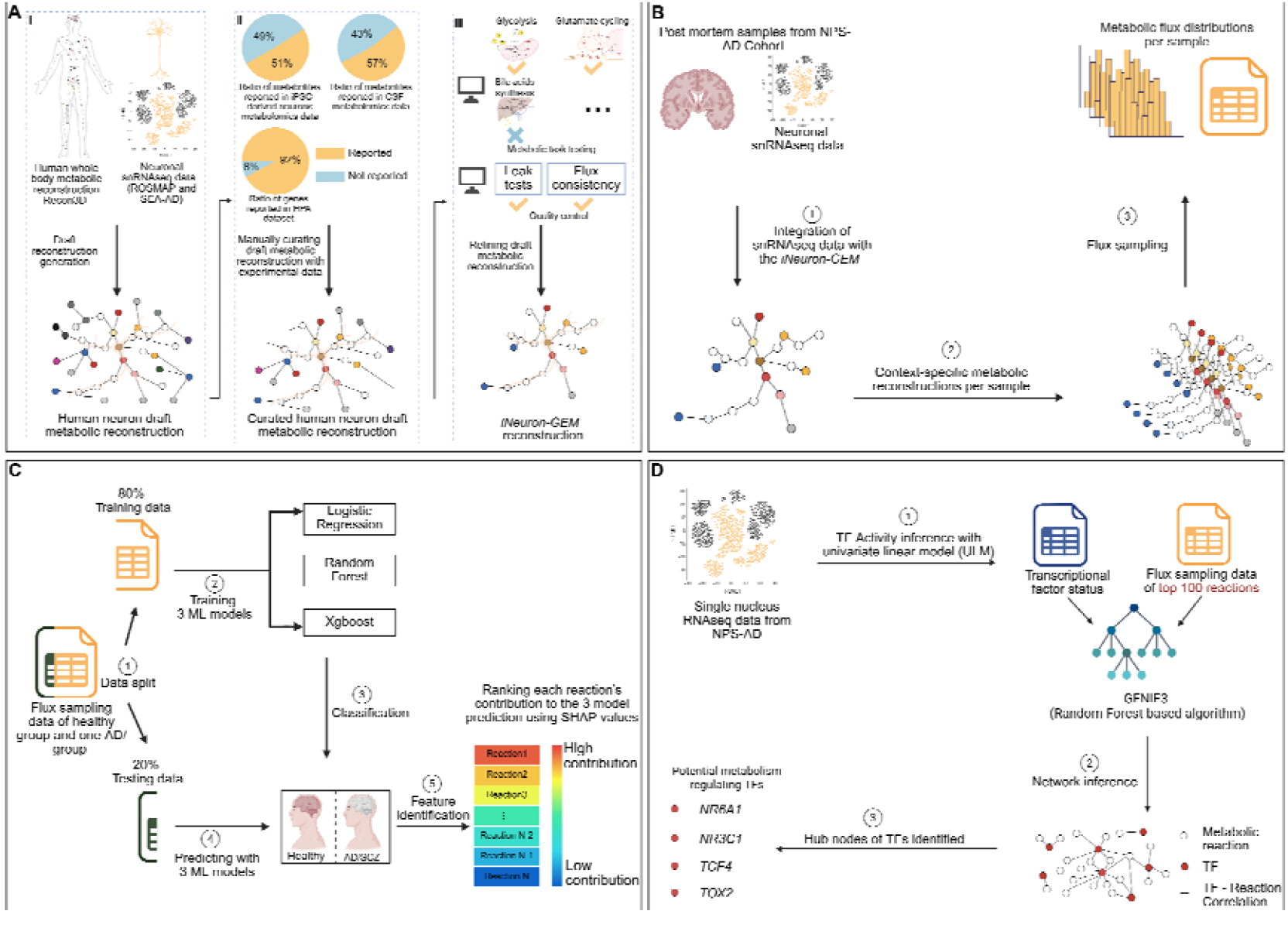
Overall analysis workflow. A) Generation of human neuron specific metabolic reconstruction, *iNeuron-GEM*; B) Integration of snRNA-seq data from the NPS-AD study with *iNeuron-GEM* and flux sampling; C) Application of machine learning (ML) on the flux sampling outputs to filter and identify important metabolic reactions; D). Prediction of potential transcription factors (TFs) that regulate predicted metabolic disruptions of neurons in AD.

### Testing cell type-specific metabolic tasks and assessing quality of metabolic network

We carried out *in silico* metabolic task testing to ensure our *iNeuron-GEM* can perform metabolic activities essential for and specific to human neurons (Figure 1A III). A total of 196 metabolic tasks for mammalian cells were collected from literature and tested in the *iNeuron-GEM* (Supplementary File 1 Table S3). 79% of tasks including metabolic processes important for human neurons, such as glycolysis, oxidative phosphorylation for ATP production, synthesis of neurotransmitters such as glutamate, glutamine and GABA, and degradation of amino acids (e.g. glycine and arginine) were feasible in our metabolic network. We also performed sanity checks by including tasks not related to neurons. Some of these tasks included synthesis of bilirubin and bile acids, and synthesis of glycogen. Our metabolic network did not pass these tests, indicating cell type-specificity. Quality control of the metabolic network^19^ was aimed to ensure that *iNeuron-GEM* does not generate ATP or produce metabolites when there are no essential metabolite inputs and also the network is free of dead end metabolites (Supplementary File 1 Table S4). All these checks ensured that the model is leak-proof and can physiologically mimic neuronal cells.

### Identifying metabolic signatures of neuronal subtypes in neurodegenerative and neuropsychiatric disorders

#### Context-specific reconstructions from snRNA-seq data

To explore metabolic alterations in neurodegenerative and neuropsychiatric disorders, we extracted snRNA-seq data of 1499 human postmortem prefrontal cortex samples from the Mount Sinai Neuropsychiatric Symptoms in AD (NPS-AD) study^17^. From each sample, expression data of both excitatory neurons (EN) and inhibitory neurons (IN) were extracted. Using the iMAT algorithm^23^, neuronal data was integrated with our *iNeuron-GEM* network to generate context-specific metabolic networks for the two neuronal subtypes (Figure 1B). We generated 2889 context-specific metabolic reconstructions and explored metabolic flux distributions in IN and EN from all samples.

#### Exploring metabolic alterations across different stages of AD and in schizophrenia

Based on the clinical diagnosis and Braak stages provided in NPS-AD, samples were grouped into six AD groups based on their assigned Braak stages and healthy controls (Figure 2A). To explore *in silico* metabolic flux distributions in each context-specific metabolic networks, we used Monte Carlo Artificially Centered Hit and Run (ACHR) flux sampling algorithm to randomly select 100 feasible solution points in its allowable solution space (Figure 1B). Outputs of flux sampling enabled an unbiased assessment of all possible flux distributions of metabolic networks from different groups^24^. Results of one-way ANOVA and Tukey’s *post hoc* test illustrated that EN had more reactions with significant flux differences between the healthy group (Braak stage 0) and other six AD groups (Braak stage 1-6) than the IN (Figure 2B). Mapping these reactions to metabolic subsystems showed that in both EN and IN, more than 50% reactions with significantly different fluxes belonged to fatty acid oxidation (average number of reactions-IN: 222; EN: 349), fatty acid synthesis (average num of reactions-IN: 70; EN: 119), cholesterol metabolism (average number of reactions-IN: 72; EN: 86), bile acid metabolism (average num of reactions-IN: 17; EN: 26) and glycolysis (average number of reactions-IN: 20; EN: 22). Numbers of these disrupted reactions were higher in EN than in IN (Figure 2C, D).

**Figure 2.**
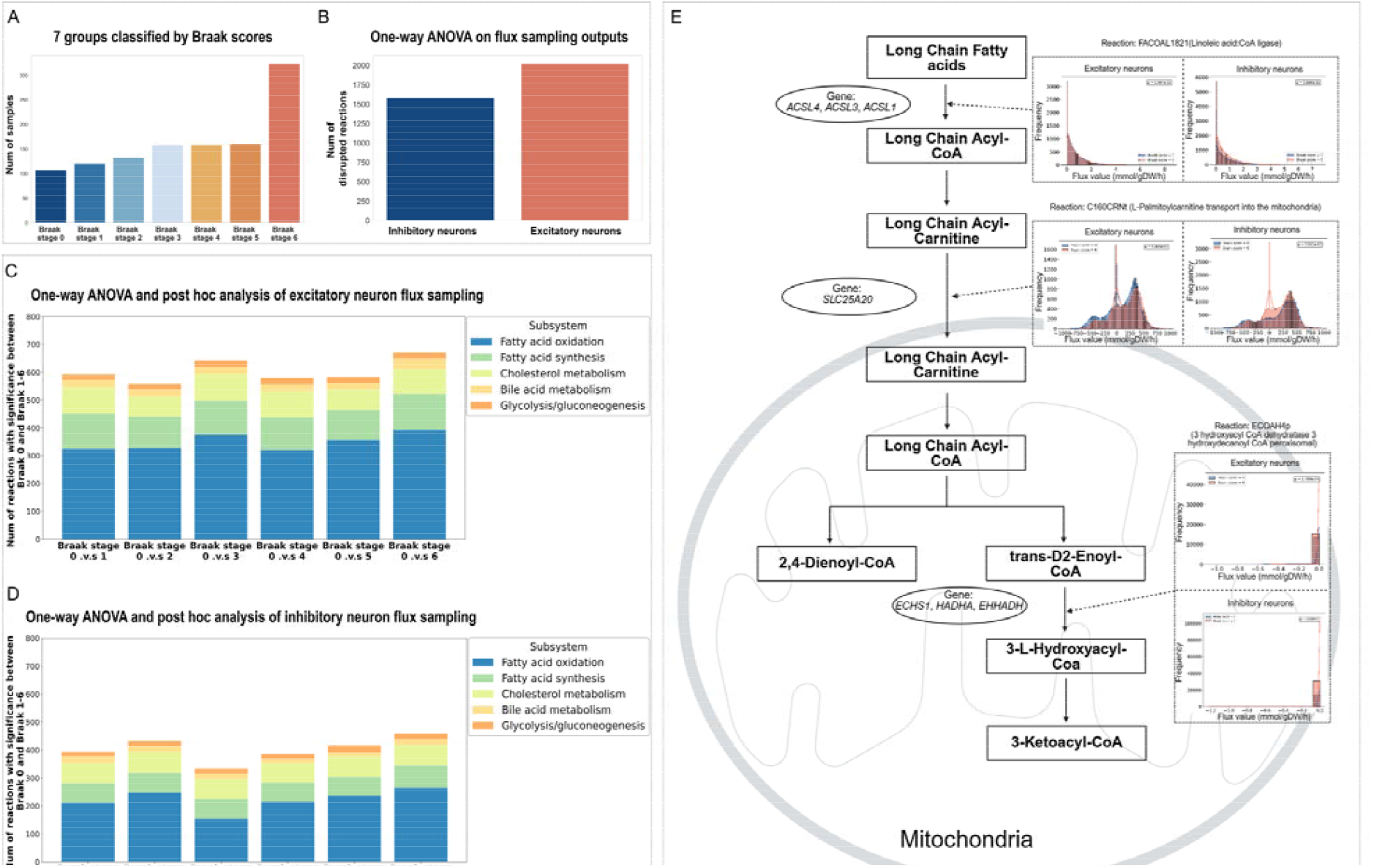
Flux balance analysis on neurons of AD. A) Bar plot illustrating the number of samples among 7 groups based on Braak stages of individuals, Braak stages 0 means healthy individuals and Braak stages 1-6 means AD patients ; B) Bar plot illustrating number of reactions with significant flux difference between healthy and AD in EN and IN (one-way ANOVA, *p*-value□<□0.05 ; C,D) Stacked bar plots illustrating number of reactions with significant flux difference between Braak stages 0 and Braak stages 1-6 in EN and IN (Tukey’s *post hoc test*, adj-p-value <0.05); E). An example illustrating the reactions belonging to long chain fatty acid oxidation and with different metabolic flux distributions between Braak stages 0 and Braak stages 1-6 in EN and IN.

In the NPS-AD data, we also extracted and stratified the samples into healthy versus SCZ group based on the diagnosis of SCZ (Figure 3A). Results illustrated that similar to AD, both EN and IN had reactions with significantly altered fluxes between healthy group and SCZ group. A higher proportion of these reactions belonged to fatty acid oxidation (num of reactions-IN: 258; EN: 407), fatty acid synthesis (num of reactions-IN: 71; EN: 142), cholesterol metabolism (num of reactions-IN: 83; EN: 82), bile acid metabolism (num of reactions-IN: 20; EN: 28) and glycolysis (num of reactions-IN: 14; EN: 19). In addition, EN had significantly more disrupted reactions in fatty acid oxidation and fatty acid synthesis than IN (Figure 3B).

**Figure 3.**
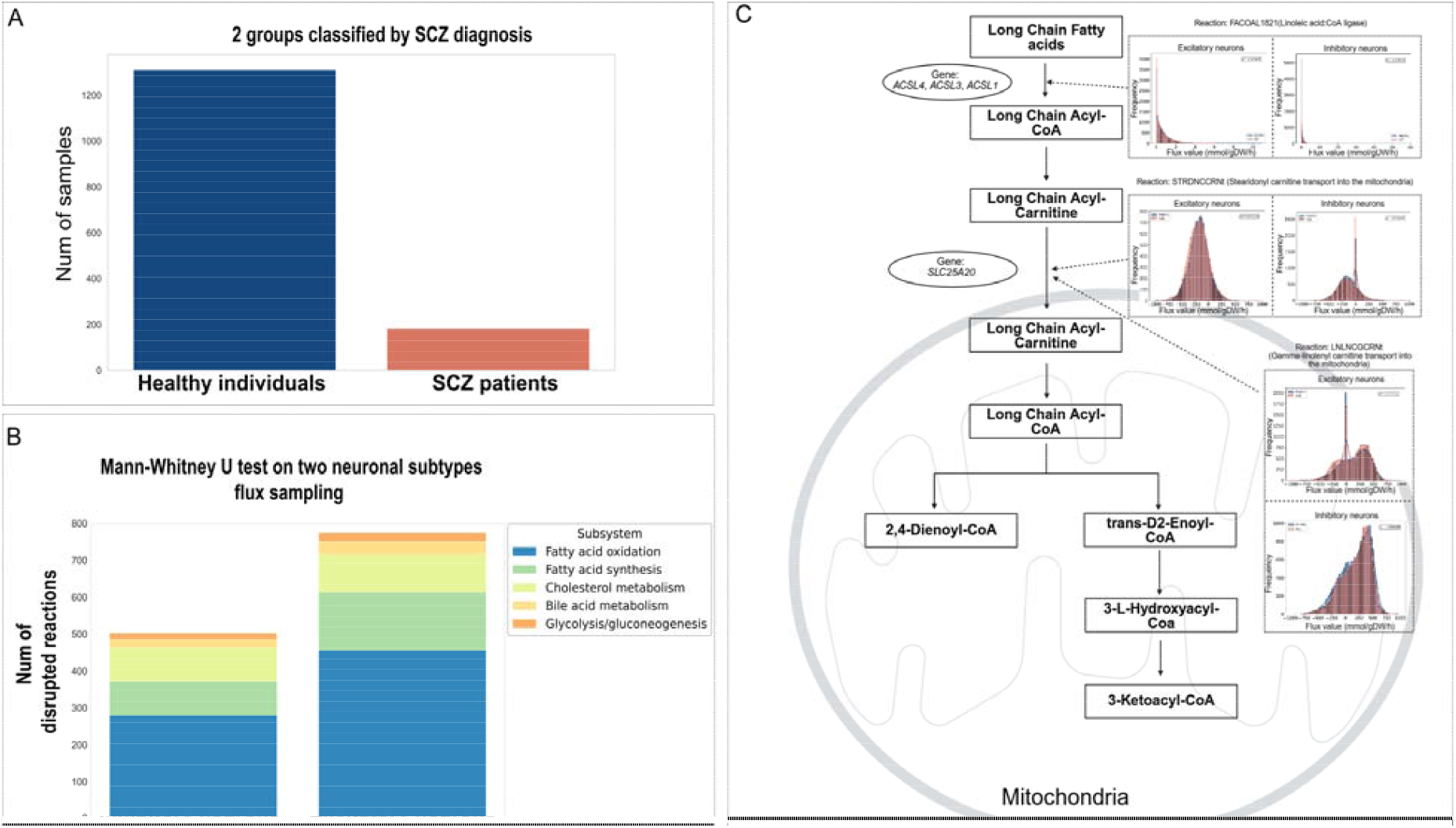
Flux balance analysis on neurons of SCZ. A) Bar plot illustrating the number of samples in the healthy group and SCZ group; B) Stacked bar plots illustrating number of reactions with significant flux difference between healthy individuals and SCZ patients in EN and IN (*student’s t-test*, p-value < 0.01); C). An example illustrating the reactions belonging to long chain fatty acid oxidation and with different metabolic flux between healthy individuals and SCZ patients in EN and IN.

#### Identification of important neuronal subtype metabolic alterations in AD and SCZ

We further investigated whether the predicted *in silico* metabolic reaction fluxes of IN and EN could differentiate samples from the healthy group from the disease groups (like Healthy versus AD groups, and healthy versus SCZ groups), using machine learning (ML) approaches. After training three ML models, (logistic regression, random forest, and XGBoost) with the flux data and running a 10-fold cross-validation, we found that based on our flux data, the three ML models can identify features between healthy individuals and AD/SCZ patients, with scores of accuracy, precision, area under the receiver operating characteristic curve (AUROC) and area under the precision recall curve (PR-AUC) of over 0.8 (Supplementary File 1 Table S5). Based on the reasonable classification performance of ML models, we then employed SHAP (SHapley Additive exPlanations) values ^25^ to determine top 100 features (Supplementary File 1 Table S6), namely reactions, that had high contributions to ML classifications(Figure 1C). Genes, metabolites and metabolic fluxes associated with identified reactions could be potential metabolic signatures of AD and SCZ.

In the AD dataset, we mapped genes corresponding to the top 100 reactions in both IN and EN across Braak stage 1-6 to a list of nominated AD targets from Agora (an database for exploration of AD evidence based on omics data) ^26^. The result illustrated that among the metabolic genes of the top 100 reactions, there were 41 genes in EN and IN that overlapped with potential AD targets. These mapped AD targets had different distributions in EN and IN across different Braak stages (Figure 4A). For example, 21 metabolic genes including *ETFA, PDHA1* and *ACO2* were identified in both EN and IN, and they were involved in TCA cycle and inositol phosphate metabolism (Figure 4C). The 20 AD targets identified only in EN, such as *DBT* and *NDUFS7*, were enriched in oxidative phosphorylation and the neurodegeneration pathway (Figure 4D). The genes identified only in IN, such as *PLPPR4* and *AADAT* (Figure 4E), were associated with lipid and amino acid metabolism. We also observed that in the later AD stages (Braak stage 5-6), EN showed more overlapping AD target genes compared to early and mid AD stages (Braak stage 1-4). In contrast, numbers of mapped AD target genes in IN were similar across Braak stage 1-6.

**Figure 4.**
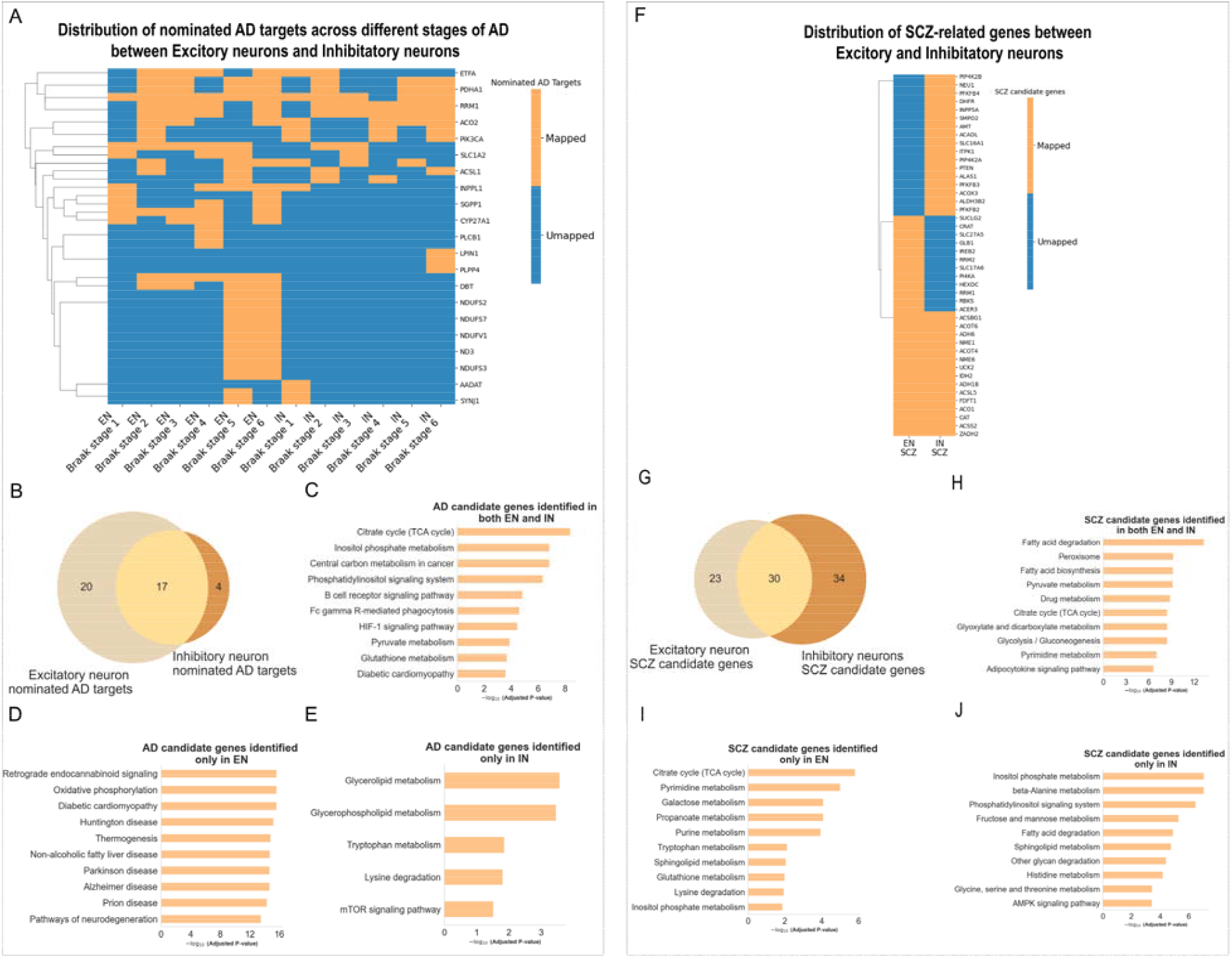
Feature filter and identification with ML. A) Heatmap illustrating the distribution of identified AD targets in EN and IN across Braak stage 1-6; B) Venn diagram comparing numbers of identified AD target genes between EN and IN ; C, D, E) Pathway enrichment on the AD targets identified in both EN and IN, and identified exclusively in EN and IN; F) Heatmap illustrating the distribution of identified SCZ targets in EN and IN between healthy and SCZ conditions; G) Venn diagram comparing numbers of identified SCZ target genes between EN and IN; H, I, J).) Pathway enrichments on the SCZ targets identified in both EN and IN, and identified exclusively in EN and IN.

In the SCZ dataset, we mapped 87 metabolic genes associated with the top 100 reactions in both IN and EN to SCZ related genes from the SZDB database (a comprehensive database for schizophrenia research) (http://szdb.org) ^27^. However, in contrast to AD, the IN of SCZ had more SCZ associated genes than EN (Figure 4B, 4G), and 34 of them were only identified in IN (Supplementary File 1 Table S6), which were involved in amino acid metabolism, inositol phosphate metabolism and phosphatidylinositol signaling system (Figure 4J). While in EN of SCZ, its 23 unique SCZ associated genes were enriched in TCA cycle, pyrimidine metabolism and propanoate metabolism (Figure 4I).

#### Exploring correlation between transcription factors and metabolic fluxes for the neuronal subtypes across different stages of AD

To explore potential mechanisms driving neuronal metabolic alterations in AD, we first inferred TF activities of EN and IN in each NPS-AD sample, using decoupleR ^28^. Correlations between 170 neuron enriched TFs activities and flux sampling output were analyzed by adapting the well-known GENIE3 method ^29^ (Figure 1D). Hub nodes of TFS in the transcription factor-metabolic flux correlation network were further identified in Cytoscape ^30^ (Figure 5B). All identified TFs showed significantly different activities between samples of healthy group and samples of AD patients from corresponding Braak stages (Figure 5A). In both EN and IN, we identified about 12-16 hub nodes of TFs across different Braak stages, some of which demonstrated distinct distributions during AD progression (Supplementary File 1 Table S6). For example, in EN, *NPAS2* was only identified in the early AD stage (Braak stages 1-2), while *NR6A1, NR3C1, TCF4* and *SOX11, PRDM10* and *MRTFA* were identified in both middle (Braak stages 3-4) and later AD stages (Braak stages 5-6). In IN, *NR6A1 was* identified as a hub node of TFs, but only in the early and mid AD stages. When comparing TFs between EN and IN, at the Braak stage 1-2, only one TF (*DACH1*) was identified in both EN and IN, while there were 7 TFs (*NR3C1, NR6A1, NRF1, RARB, RORA, SETBP1* and *SOX11*) commonly identified in both neuron subtypes at Braak stage 3-4, and 5 TFs (*GTF2IRD1, MRTFA, PBX1, TCF4, TOX2*) at Braak stage 5-6 (Figure 5C).

**Figure 5.**
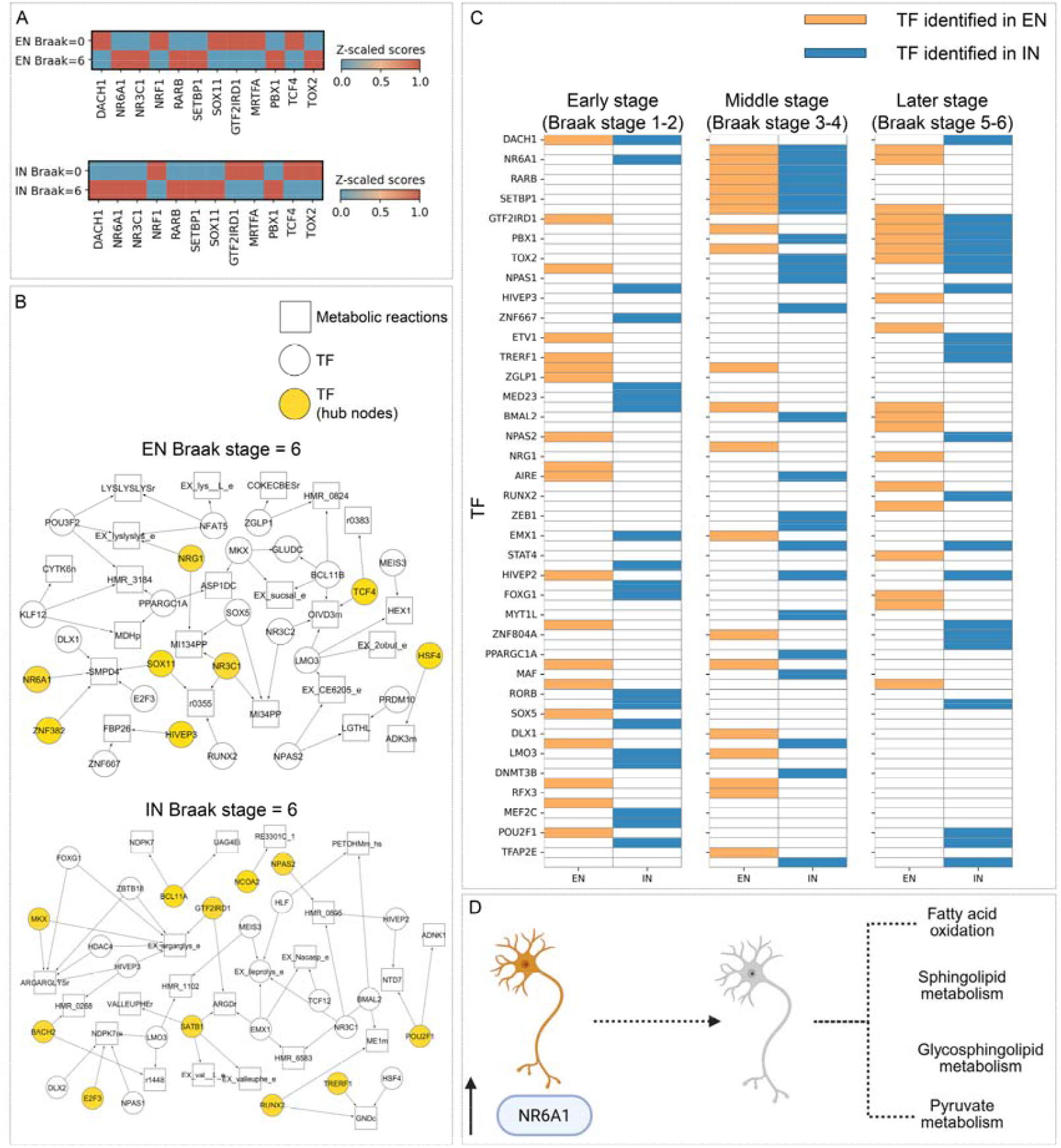
Metabolism-related TF prediction. A) Heatmap comparing normalized TF activity scores of 12 TFs between Braak stage 0 and Braak stage 6 in EN and IN; B) Example of network illustrating correlation between TFs and metabolic reaction fluxes in Braak stage 6 in EN and IN; C) Heatmap illustrating distributions of identified hub node TFs among different stages of AD in EN and IN; D) Diagram illustrating potential correlation between NR6A1 and disruption of lipid and energy metabolism in neurons.

Results also illustrated that these identified TFs may regulate different metabolic activities between EN and IN (Supplementary File 1 Table S7). For instance, in EN, *NR6A1* had a high correlation with the simulated *in silico* metabolic fluxes of reactions belonging to sphingolipid metabolism (reactions id: SMPD4), fatty acid oxidation (reactions id: HMR_3184), glycosphingolipid metabolism (reactions id: HMR_0824) androgen and estrogen synthesis and metabolism (reactions ids: HMR_1976 and RE3108C) (Figure 5D); In comparison, *NR6A1* in IN had a high correlation with metabolic reactions of fatty acid oxidation (reactions ids: CSNATr, HMR_0386 and r1448), pyruvate metabolism (reactions ids: ALCD22_D and PPDOy) and extracellular metabolite exchange (reactions ids: EX_val L_e and EX_anth_e).

## Discussion

In this study we integrated *in silico* metabolic modeling and machine learning (ML) to systematically investigate human neuronal subtype-specific metabolic alterations and their potential regulatory mechanisms in AD. The resulting *iNeuron-GEM* represents, to our knowledge, the first manually curated, genome-scale metabolic network of human neurons. Its reconstruction involved a) single nucleus RNA sequencing (snRNA-seq) data of human postmortem prefrontal cortex samples from two well-characterized AD cohort studies, namely ROSMAP^12^ and SEA-AD^31^; b) metabolites detected in the untargeted metabolomics analyses of human iPSC-derived neurons and human cerebrospinal fluid (CSF) (Table 1); c) neuronal genes with a non-zero nTPM values from the Human Protein Atlas^20^. These datasets ensured that the genes, metabolites and reactions in *iNeuron-GEM* network accurately captured neuron-specific metabolic functions. Model validation against 196 *in silico* metabolic tasks confirmed its ability to perform essential cellular and neuronal processes, including glycolysis, oxidative phosphorylation, the citric acid cycle, and neurotransmitter synthesis and transport (Supplementary File 1 Table S3). Flux-consistency checks further confirmed the absence of futile loops ^19^. The final iNeuron-GEM therefore provides a robust platform to simulate steady-state flux distributions underlying human neuronal metabolism.

By integrating snRNA-seq data from excitatory (EN) and inhibitory (IN) neurons in the NPS-AD cohort ^17^ with our *iNeuron-GEM* network, we identified distinct and overlapping metabolic alterations across AD Braak stages and in schizophrenia (SCZ). We identified that EN exhibited more metabolic alterations than IN across Braak stages 1-6. Previous studies have shown that inhibitory neurons are lost early in AD progression, contributing to excitatory/inhibitory (E/I) imbalance^32^. IN terminals sustained higher levels of ATP indicating differential dependance on mitochondria metabolism^33^. Our analysis revealed that the majority of disrupted reactions in both EN and IN involved fatty acid and cholesterol metabolism (Fig. 2E). Although astrocytes are traditionally considered the primary site of fatty acid oxidation ^34^, our reconstruction highlighted that neuronal fatty acid metabolism occupies a substantial portion of the human neuron metabolic network (761 of 3,900 reactions across 87 subsystems; Supplementary Table S1a). This aligns with lipidomics studies detecting abundant lipid species, such as phosphatidylcholine (PC) and phosphatidylethanolamine (PE), in human iPSC-derived neurons^6^.

Among the top 100 reactions identified by ML, several fatty acid oxidation reactions— including PTDCACRNt (pentadecanoate transport into mitochondria) and DMNONCRNt (4,8-dimethylnonanoyl carnitine transport)—were key features distinguishing AD and SCZ from healthy states (Figs. 2E, 3C). Although neuronal fatty acid metabolism in AD has been understudied, recent evidence suggests that fatty acids can serve as an alternative energy source for synaptic activity via β-oxidation^35^. As β-oxidation occurs within mitochondria, the disruptions observed in lipid and fatty acid metabolism in both AD and SCZ neurons likely reflect mitochondrial dysfunction^36^. In parallel, consistent alterations were also observed in glycolysis, the citric acid cycle, and pyruvate metabolism, pathways central to neuronal energy homeostasis and previously implicated in aging and neurodegeneration^37,38^.

Mapping genes associated with the top disrupted reactions against curated AD (Agora database) and SCZ (SZDB) gene lists revealed that our framework identified known metabolic genes implicated in these disorders. The differential distribution of AD-associated targets between EN and IN across Braak stages suggests subtype-specific vulnerability, with EN showing a higher number of disrupted AD targets, that potentially reflects later-stage degeneration following early IN loss^39^. Enrichment of 21 shared AD target genes in both EN and IN revealed strong involvement in lipid and energy metabolism and the HIF-1 (hypoxia-inducible factor 1) signaling pathway. The HIF-1 pathway mediates hypoxic responses and has been proposed as a therapeutic target for AD^40^. Recent studies also demonstrated that disrupted HIF-1α homeostasis was correlated with increased oxidative stress, hyperglycemia, insulin resistance, and inflammation in both AD and type 2 diabetes mellitus^41^. Our identification of convergent lipid, energy, and HIF-1 pathway disruptions supports the view that impaired bioenergetic metabolism is a unifying hallmark across AD neuronal subtypes.

In SCZ, 87 metabolic genes were mapped to SZDB, with IN showing a higher number of SCZ-associated metabolic genes than EN despite fewer total reaction-level changes (Fig. 4G). Three genes, namely DLD, ACO2, and IDH2, were consistently identified across both neuronal subtypes and diseases. ACO2 encodes mitochondrial aconitase 2, a citric acid cycle enzyme sensitive to oxidative stress^42^ and has been increasingly emphasized in neurodegenerative disorders as a key metabolic factor^43^. Although studies on *ACO2* in SCZ were limited, a recent study based on metabolomics data derived from SCZ patients reported that the level of isocitrate in first-episode schizophrenia patients were up-regulated^44^. IDH2 catalyzes isocitrate oxidation to α-ketoglutarate, generating NADPH to protect against oxidative damage, was also highlighted as novel diagnostic biomarker panel for SCZ ^45^. Both genes have been linked to disrupted mitochondrial metabolism in neurodegenerative and psychiatric disorders, suggesting shared vulnerabilities in neuronal mitochondrial function across AD and SCZ.

Transcriptional regulation analysis identified 13 transcription factors (TFs) whose activity scores correlated strongly with *in silico* flux changes across Braak stages (Supplementary File 1 Tables S7–S8). The number of shared TFs increased from early (Braak 1–2) to late (Braak 5–6) stages, implying progressive convergence of transcriptional dysregulation. Among these, NR6A1 emerged as a key regulatory hub, correlating with sphingolipid, fatty acid, and glycosphingolipid metabolism. NR6A1, an orphan nuclear receptor, has been implicated in embryonic neuronal differentiation and lipid metabolism regulation via the mTORC1 pathway ^46^. In mouse hippocampal neurons, knockdown of *NR6A1* was reported to protect mice from depression by mediating the cyclic adenosine monophosphate (cAMP)-response element binding protein (CREB)-brain-derived neurotrophic factor (BDNF) pathway ^47^. Another recent study on HT-22 neuronal cells further demonstrated that *NR6A1* silencing had protective effects against neuronal mitochondrial disruptions ^48^. Although direct evidence linking NR6A1 to lipid dysregulation in AD is lacking, our findings nominate it as a promising target for mechanistic investigation.

In summary, our study introduces the first curated human neuron–specific GEM and demonstrates its utility in dissecting neuronal subtype-specific metabolic alterations in AD and SCZ. By integrating *in silico* flux modeling with ML, we identified convergent disruptions in lipid and energy metabolism, recapitulated known disease-associated genes, and predicted novel TF regulators such as NR6A1. While iNeuron-GEM provides valuable insights, several limitations remain. The model simulates steady-state fluxes and does not capture dynamic enzyme kinetics or extracellular metabolite concentrations ^50^, and it does not directly quantify absolute metabolite levels. Future refinements incorporating experimental metabolomics and fluxomics data will enable dynamic simulation and improved accuracy. Such integrative frameworks will advance our understanding of neuronal metabolic reprogramming and facilitate the identification of therapeutic targets for neurodegenerative and neuropsychiatric diseases.

## Method

### Collecting and processing single-nucleus RNA sequencing data of human neurons

From the AD Knowledge Portal, we collected single-nucleus RNA sequencing data from the prefrontal cortex of postmortem samples from two AD cohorts - The Religious Orders Study and Memory and Aging Project (ROSMAP) Study^12^ and The Seattle Alzheimer’s Disease Brain Atlas (SEA-AD) Study^31^. A total of 1034 samples of unique individuals over 70 years old were collected (Supplementary File 1 Table 1). In each sample, we imputed and extracted expression data of both excitatory and inhibitory neurons using the Seurat toolkit (http://www.satijalab.org/seurat)^51^ and cell markers of human neurons.

### Draft reconstruction of human neurons metabolic network

With the expression data of human neurons, we calculated ubiquity scores ^52^ of genes across all 1034 samples. Ubiquity scores were used to decide the confidence that a gene exists in the human neuron. Then according to the workflow of CORDA (https://github.com/resendislab/corda) ^53^, an established tool for metabolic network reconstruction, we used the human whole body metabolic reconstruction—Recon3D ^19^ and the ubiquity score as inputs to run CORDA, whose output was a draft metabolic reconstruction of human neurons.

### Curation and refinement of human neuron draft reconstruction

The draft reconstruction of human neurons was tested with 229 metabolic tasks (Supplementary File 1 Table S2) *in silico*. To increase the specificity of the human neuron reconstruction, we ensured that it could achieve a set of metabolic tasks reported to be essential to human neurons. For example, the reconstruction must be feasible to produce ATP from glucose both with and without oxygen, which simulated the glycolysis and oxidative phosphorylation process in the human neurons. Then, we further curated the reconstruction with experiment data of both metabolomics and transcriptomics. The untargeted metabolomic data of human iPSC-derived neurons and CSF were collected from publications and public databases (Table 1). All collected metabolites were standardized with HMDB ID and then mapped to the virtual metabolic human database ^42^. Based on the collected metabolome and transcriptome data, we further curated the metabolites, genes and metabolic reactions in the reconstruction to ensure that most of them are consistent with experimental data. For example, if most of the metabolites and genes associated with one reaction were not supported by experimental data, then this reaction would be removed from the reconstruction if it did not cause large range of inconsistency in the metabolic network.

### Testing basic properties of reconstruction

We checked the quality of the curated human neuron reconstruction with a series of tests using the COBRA Toolbox. With these tests, e.g. the leak and sanity test^54^, we ensured that in the *in silico* simulation, (1) the reconstruction did not produce any metabolites when its exchange and sink reactions were closed; (2) the reconstruction did not produce ATP from water or oxygen; (3) the reconstruction did not have dead ends in the network. After passing the above tests, we future used MEMOTE^55^, an established the genome-scale metabolic model test suite, to ensure the quality of the human neuron metabolic reconstruction.

### Context-specific metabolic reconstruction derived from patients with neurodegenerative and neuropsychiatric disorders and simulation

SnRNA-seq data from the dorsolateral prefrontal cortex of 1494 postmortem samples were collected from the NPS-AD study^17^. It contained the diagnosis information of AD and SCZ of individuals. With Seurat, we ran imputation on each sample and extracted expression data of excitatory (EN) and inhibitory (IN) neurons, separately. Then gene expression data of excitatory and inhibitory neurons were normalized into counts per million (CPM). We used iMAT^23^, a well-known tool for building context-specific metabolic reconstruction, to integrate CPM of each sample with human neuron metabolic reconstruction, to build 2988 context-specific metabolic reconstructions of both excitatory and inhibitory neurons. To simulate metabolic flux distributions in neurons from different groups, we employed the ACHR flux sampling algorithm from the COBRApy^56^ to randomly sample feasible steady-state fluxes for 1000 times in the allowable solution spaces among context-specific metabolic reconstructions. *In silico* metabolic reaction flux distributions across AD and SCZ stages between excitatory and inhibitory neurons were compared based on results of flux sampling. Reactions with significantly different fluxes values between the healthy group and Braak stage 1-6 groups in AD were compared and analyzed with the One-way ANOVA, post hoc test (TukeyHSD test). Reactions with significantly different fluxes values between the healthy group and disease group in SCZ were compared and identified by Mann– Whitney U test distributions of reaction fluxes in both AD and SCZ were compared with Kolmogorov–Smirnov test.

### Construction of 3 machine leaning models and SHAPE values of important reactions

Flux sampling data of all samples were grouped by the diagnosis of AD and SCZ. For AD, based on the Braak score, all samples were divided into 7 groups. Samples with Braak score 0 were healthy group, and Samples with Braak score 1-6 were AD groups; while for SCZ, all samples were divided into healthy group and SCZ group. To compare the metabolic difference between the healthy group and other disease groups separately, 80% samples in each group randomly were selected as the training dataset and the rest 20% samples were selected as the testing dataset. Then 3 machine learning models including logistic regression, random forest and xgboost, were trained with training datasets of the healthy group and one individual diseases group, respectively. The three types of machine learning models served as classifiers to make classification between healthy samples and disease (AD/SCZ) groups. Hyperparameters of each machine learning model were tuned with hyperopt (v0.2.7), a python library for hyperparameter optimization using Bayesian optimization strategy ^57^. Performance of each model was evaluated by:

1. Accuray
2. Precision
3. Area under the receiver operating characteristic curve (AUROC)
4. Area under the precision recall (PR) curve

After training and hyperparameter tuning, the importance of features, which are metabolic reactions, in each model were determined by calculating SHAP (SHapley Additive exPlanations) values^25^ in them. SHAP values of each feature in each model represented degree of contributions of that feature to models’ final classification performance. For each comparison between the heathy group and one disease group, average SHAP values of each feature were calculated and the top 50 metabolic reactions with the highest average SHAP values were selected as the important features to further explore and explain metabolic differences between the healthy group and different disease groups. Furthermore, genes associated with the selected reactions were mapped with nominated AD targets from Agora database(https://agora.adknowledgeportal.org/)^26^ and SCZ candidate genes from SZGR2 database(https://bioinfo.uth.edu/SZGR/)^27^.

### Correlation between transcriptional factor status and metabolic reactions

Transcriptional factor status in excitatory and inhibitory neurons of each sample was inferred with decoupleR ^28^, following its official vignette. Then we adapted the GENIE3 algorithm(https://github.com/vahuynh/GENIE3/tree/master), which was based on regression trees to explore gene regulatory networks from expression data^58^, to infer correlations between transcriptional factor status and metabolic reaction flux distribution. For each group, including healthy groups (AD and SCZ), AD groups (Braak stage 1-6) and SCZ group, status of all transcriptional factors (features) and flux sampling data of each metabolic reaction (labels) were used to train a random forest regressor. Then feature importances of each random forest regressor were extracted. Scores of links or edges between transcriptional factors and metabolic reactions were decided by the feature importances of transcriptional factors to each metabolic reaction. Furthermore, Cytoscape (v3.10.3)^30^ were used to identify hub nodes in the network consisted with of transcriptional factors to all metabolic reactions in each group.

## Data availability

The snRNAseq used to generate iNeuron-GEM reconstruction from the ROSMAP and SEA-AD studies are available in Synapse (https://www.synapse.org) (Synapse ID syn23650894), and Seattle Alzheimer’s Disease Brain Cell Atlas (https://portal.brain-map.org/explore/seattle-alzheimers-disease), respectively. The snRNAseq data used for the case study from the NPS-AD study is available in Synapse (ID:syn60543127). The data of neuronal cell type-specific proteins was extracted from Human Protein Atlas (https://www.proteinatlas.org).

## Code availability

Codes used for iNeuron-GEM generation, data integration and ML analysis are available in the following GitHub repository: https://github.com/BaloniLab/iNeuron-GEM

## Acknowledgements

This study was funded by grants from the National Institute of Environmental Health Sciences (NIEHS) (ES010563, ES031401 (A.B.B., H.K), AG080917 (A.B.B., A.M.T.), ES010563 (A.B.B., P.B.)). A.M.T. was additionally supported by a K99/R00 Pathway to Independence Award (K99 ES036290). The authors thank the participants and investigators of the ROSMAP, SEA-AD, and NPS-AD cohorts for making their data available in AD Knowledge Portal and acknowledge the Purdue University School of Health Sciences and EMBRIO for institutional support.

